# Interplay between metrical and semantic processing in French: an N400 study

**DOI:** 10.1101/738930

**Authors:** Noémie te Rietmolen, Radouane El Yagoubi, Corine Astésano

## Abstract

French accentuation is held to belong to the level of the phrase. Consequently French is considered ‘a language without accent’ with speakers that are ‘deaf to stress’. Recent ERP-studies investigating the French initial accent (IA) however demonstrate listeners to not only discriminate between different stress patterns, but also expect words to be marked with IA early in the process of speech comprehension. Still, as words were presented in isolation, it remains unclear whether the preference applied to the lexical or to the phrasal level. In the current ERP-study, we address this ambiguity and manipulate IA on words embedded in a sentence. Furthermore, we orthogonally manipulate semantic congruity to investigate the interplay between accentuation and later speech processing stages. Results reveal an early fronto-centrally located negative deflection when words are presented without IA, indicating a general dispreference for words presented without IA. Additionally, we found an effect of semantic congruity in the centro-parietal region (the traditional region for N400), which was bigger for words without IA than for words with IA. Furthermore, we observed an interaction between metrical structure and semantic congruity such that ±IA continued to modulate N400 amplitude fronto-centrally, but only in the sentences that were semantically incongruent. The results indicate that presenting word without initial accent hinders semantic conflict resolution. This interpretation is supported by the behavioral data which show that participants were slower and made more errors words had been presented without IA. As participants attended to the semantic content of the sentences, the finding underlines the automaticity of stress processing and indicates that IA may be encoded at a lexical level where it facilitates semantic processing.

## 1 Introduction

Prosody has an important role in speech comprehension; where in written form, language is structured by white spaces and punctuation marks, spoken language is organized through intonation, accentuation, and rhythm. Especially metrical structures have long been considered crucial in the segmentation of speech. With no clear separation between words in the speech signal, the metrical segmentation strategy (MSS) proposes that listeners rely on their languages’ metrical pattern to identify word boundaries (Cutler & Norris, 1988; Cutler, 1990). Indeed, in stress-based languages, such as English or Dutch, wherein stress is part of the lexical entry, accents provide reliable cues to lexical boundaries. And also in French, a language often described to be syllable-based due to the fairly homogeneous metrical weight on syllables, prosodic structure has been found to guide speech segmentation (Welby, 2007; Spinelli et al., 2010; Banel & Bacri, 1994; Bagou et al., 2002; Christophe et al., 2004, among others). However, in these studies, segmentation was not considered lexical but presumed phrasal, i.e. listeners are assumed to adopt a *prosodic segmentation strategy* in which intonational and accentual patterns function to segment *prosodic groups* (i.e. level of the accentual phrase, AP; Jun & Fougeron, 2000) from the speech signal (Wauquier-Gravelines, 1999). This view stems from traditional descriptions of French as ‘a boundary language’ (Vaissière, 1991) or ‘a language without accent’ (Rossi, 1980) according to which stress, because it is not lexically distinctive in French and because its surface realization is acoustically merged with intonational boundaries, has no clear metrical value.

Two group-level accents are generally recognized in French, the primary final accent (FA) and the secondary initial accent (IA). FA is the compulsory stress and falls on the last syllable of the last word of AP where it marks the right prosodic constituent boundary. Because FA relies on largely the same acoustic-phonetic parameter as intonation in French, local prominences near phrase boundaries will blend with the intonation contour so that their phonetic parameters are spread and diluted over adjacent syllables (e.g. Rossi, 1980; Fónagy, 1980). Moreover, when a word is embedded into a phrase, primary accents within the phrase may be phonetically reduced, or de-accented (e.g. Di Cristo, 1999; Astésano, 2016), to favor a more prominent marking of the phrase boundary (hence the label *boundary language* for French, Vaissière, 1991). For instance, in ‘jolie fille’ the primary accent on the phrase internal word (‘jo<u>lie</u>’) may be reduced such that the group boundary is more pronounced (‘jolie <u>fille</u>’) (Delattre, 1966; Rossi, 1980). It is important to realize, however, that ‘de-accentuation’ does not mean that the accent is deleted and disappears completely. Instead, the accent is reduced to various degrees depending on rhythmic, contextual and pragmatic circumstances. This means both that 1) a trace of the local prominence survives and that 2) de-accentuation does not exclusively serve a clear marking of phrasal boundaries. In the above example, for instance, the de-accentuation also helped dodge a stress clash when the primary French accent (FA) located on the last syllable of ‘jo<u>lie</u>’ was followed by the monosyllabic ‘<u>fille</u>’ (also carrying primary stress). Indeed, the occurrence of two consecutive stressed syllables is universally disfavored and may be avoided by restructuring the surface realization of the underlying prosodic representations (Liberman & Prince, 1977; Nespor & Vogel, 1983).

For instance, in French, de-accenting FA to evade stress clashes may lead to the first syllable of the phrase being accented instead, the initial accent (IA, ‘<u>jo</u>lie <u>fille</u>’). This is one of the reasons for which the initial accent is interpreted as the optional and secondary accent in French. The initial accent is, however, not exclusively a result of the stress clash resolution; IA also serves a rhythmic balancing function to break long stretches of unaccented syllables, again contributing to its status as a secondary accent. Further, the accent is often confused with the emphatic accent. That is, the initial stress may also be expressive, pragmatically contrasting sentence meaning with an accentual emphasis. Finally, as in the example above, the initial accent may mark the left boundary of AP and help group the words into a cohesive union (Di Cristo, 1999; Astésano, 2016). That is, the union of IA and FA, called *an accentual arch* (Fónagy, 1980), presents a bipolar stress template which underlies AP and groups the words it contains (see also Rolland & Lœvenbruck, 2002). Thus, IA is associated with a number of different functions, but these functions remain post-lexical in nature so that the phonological status of IA and its functions in lexical processing are still unclear.

In the current ERP-study, we investigate the functional role of IA in word level processing. We align to Di Cristo’s metrical model of French which proposes both FA and IA to be phonologically encoded in (latent) cognitive stress templates underlying the representations of words (Di Cristo, 2000). According to the model, words are then marked with metrically strong syllables at both left and right lexical boundaries that can readily notify listeners on when to initiate lexical access. The model therefore provides a valuable theoretical context to speech segmentation in French. Indeed, studies showing IA to play an important role in the marking of lexical structure and speech segmentation are accumulating. For instance, a series of perception studies has found IA to be a more reliable cue to word boundaries than FA and to be perceived as more prominent at both phrasal and lexical levels (Astésano et al., 2007, 2012; Garnier et al., 2016; Garnier, 2018). Further, the initial accent is perceived even when its acoustic parameters are reduced (Jankowski et al., 1999), or when its pitch rise peaks further along in the word (e.g. Astésano et al., 2012), indicating a strong metrical expectation for the accent. These results prompted a recent paper to revisit the secondary and optional nature of IA and suggest IA carries a metrical strength that is at least equal to that of FA, both accents working together in the marking of the lexical word (Astésano & Bertrand, 2016).

Recent neuroimaging studies corroborate this idea and underline the role of IA in French word processing. When presenting words with or without IA in an oddball study, Aguilera and colleagues obtained a larger MisMatch Negativity components (MMN) when the oddball had been presented without IA than when the oddball was presented with IA (Aguilera et al., 2014). Such an asymmetry between MMNs indicates that IA is encoded in long-term memory and part of the expected stress template. Following up on these results, IA was manipulated in a lexical decision task wherein trisyllabic nouns and pseudowords were presented with or without IA (te Rietmolen et al., 2016). Omitting IA resulted in a processing cost during stress extraction as reflected by a more ample N325 (Böcker et al., 1999) regardless of lexical condition, which demonstrates both the automaticity of stress extraction and an expectation for words to be marked with IA in the pre-lexical stage of speech processing (see also Böcker et al., 1999, for a similar interpretation of the N325 in an investigation of stress processing in Dutch, a language with obligatory and distinctive stress).

However, the two ERP studies above also presented some ambiguities. Firstly, the results of the lexical decision study (te Rietmolen et al., 2016) suggested that their metrical manipulations elicited an N400, as there appeared to be a negativity in the latency range typically associated with the N400. This would indicate a role for IA in lexico-semantic processing, and so a function in word level analysis. The authors were however cautious to interpret this negativity as an N400, because words were presented in isolation (i.e. without semantic context), while the N400 is more typically elicited in paradigms such as the semantic priming paradigm (wherein a target-word directly follows a word or image to which it is semantically related or not), or the semantic anomaly paradigm (wherein sentences are presented with a target-word that is semantically congruent or incongruent within the sentence context). Secondly, presenting words in isolation had, as additional consequence, that IA was always in utterance initial position. Indeed, both in Aguilera et al. (2014) and in te Rietmolen et al. (2016) words had been presented as independent utterances, so that listeners may have processed them as individual accentual phrases. Hence, it can not be ruled out that the templates — and the processing cost when IA was omitted— applied to the phrase level instead of the level of the lexical word.

In the current N400-study we sought to elucidate these ambiguities and manipulated IA on words positioned within a sentence. Additionally, we manipulated the semantic congruity of the sentences, allowing us to investigate whether IA also affects the lexico-semantic processing stages in speech comprehension.

The N400 is a centro-parietally located negativity that peaks around 400 ms after the detection of a semantic discrepancy. The negativity is considered an adept indicator of obstructed speech comprehension, with amplitude modulations or delayed latencies revealing difficulties in speech processing. Still, the precise nature of the N400 remains a topic of considerable debate. That is, it is unclear, whether N400 modulations are restricted to semantic information or whether the N400 can additionally be modulated by mismatching phonological information, such as metrical patterns. One commonly held belief on the nature of the N400, is that it results from hindered contextual integration (van den Brink et al., 2001; Brown & Hagoort, 1993). In this view, the N400 indicates difficulties in the post-lexical stage of speech comprehension, i.e. the stage after initial pre-lexical activation and lexical access have been completed, and is unlikely to be influenced by phonological processes. Another stance, however, considers the N400 to reflect the degree of lexical pre-activation. In this view, higher levels of pre-activation (as a results of, for instance, supporting prior semantic information or word frequency) facilitate lexical access and reduce N400 amplitude (Kutas & Hillyard, 1980; Kutas & Federmeier, 2011). This stance then takes the N400 to reflect predictive, anticipatory processes that need not exclusively be of semantic nature, but can be phonological as well (DeLong et al., 2005; Lau et al., 2008).^1^

Indeed, a number of studies have shown misguided phonological expectations in healthy subjects (e.g. Praamstra & Stegeman, 1993; Dumay et al., 2001, 2002; DeLong et al., 2005) or impaired phonological analysis in patients (Robson et al., 2017) to interfere with subsequent semantic evaluation and modulate the N400. Furthermore, metrical information has also been found to interplay with lexico-semantic processing (e.g. Magne et al., 2007; Rothermich et al., 2010; Marie et al., 2011; Rothermich et al., 2012; Bohn et al., 2013). For instance, in a series of studies, Rothermich and colleagues manipulated the metrical regularity in German jabberwocky (Rothermich et al., 2010) and semantically anomalous sentences (Rothermich et al., 2012; Rothermich & Kotz, 2013) by presenting words either with a metrically regular or irregular beat and showed metrical regularity to facilitate semantic ambiguity resolution, as indicated by a modulated and earlier N400, which, unlike its usual centro-parietal distribution, appeared to be more frontally located. The authors relate their findings to theories of predictive coding and suggest metrically predictable stress to provide a metrical framework to which brain oscillations can align in an effort to optimize speech comprehension (*cf.* Pitt & Samuel, 1990). That is, by presenting speech with a regular (i.e. predictable) underlying beat, listeners were able to a priori direct their attention from one stressed syllable to the next (in their words) “island of reliability”, which in turn facilitated semantic processing. Note that Böcker et al. (1999) had a likewise interpretation of the N325 (which indeed displayed a similar latency and spatial distribution as the negativity reported in Rothermich et al. 2010, 2012) as they considered the N325 to reflect the interface of automated acoustic processing and controlled, top-down metrical processing in the analysis of speech. They argued that the N325 potentially indexes difficulties in processes that are involved in pre-lexical speech segmentation and the initiation of lexical access on the basis of rhythm and metrical stress. In that view, the N325 directly measures the role of metrical stress in speech processing as proposed in MSS (Cutler & Norris, 1988, see also the Attentional Bounce Hypothesis, Pitt & Samuel 1990). In fact, more recent work has, in a similar vein, asserted the earlier and more frontal N400 to index online speech segmentation, although in that work the frontal negativity was linked to novel word-form to conceptual knowledge mapping in parallel (e.g. Cunillera et al., 2009; Dittinger et al., 2017; François et al., 2017). So while the frontal N400 (or N325) is not yet fully understood, and has led to slightly different views as to what it precisely represents, there seems to be some common ground with phonological/metrical expectancy influencing semantic processing. That is, metrical structure helps listeners to a priori guide their attention towards stressed syllables (i.e. perceptually stable and prominent syllables located near word onsets), which cue listeners on when to segment speech and initiate their search in the mental lexicon, in turn facilitating access to meaning.

In French, a previous ERP study investigating the relationship between metrical structure and late speech processing, also found metrical violations to obstruct semantic processing (Astésano et al., 2004; Magne et al., 2007). In the study, participants listened to sentences in which semantic and/or metrical congruity was manipulated. Semantic congruity was manipulated by presenting sentences in which the last word was incoherent with the semantic context of the sentence, while metrical congruity was manipulated by lengthening the medial syllable of the last word, an illegal stress pattern in French. Furthermore, listeners completed two different tasks, one in which they attended semantic congruity, and one in which they judged metrical congruity. This allowed Magne and colleagues to determine whether metrical and/or semantic processing proceeds automatically or depends on the direction of attention. Behavioral results showed listeners to make more errors when either meter or semantics was incongruent. Furthermore, listeners made the most errors when meter was incongruent, but semantics was congruent, indicating that metric incongruities disrupt semantic processing. This interpretation was corroborated by their results from the ERP data. Not only did Magne and colleagues obtain a larger N400 to metrically incongruous words than to metrically congruous words in the metric task, but, interestingly, the metrical violation resulted in an increased N400, also in the semantic task (i.e. independent from attention), and even when the sentences were semantically congruent (see also Astésano et al., 2004). These results indicate that accentual patterns, also in French, affect the later stages of speech comprehension, during which access to meaning and semantic integration takes place.

However, in the study of Magne et al. (2007), the processing cost resulted from presenting an illegal stress pattern, with metrical weight on the medial syllable, and it remains unclear whether semantic processing also suffers when words are presented with metrical structures that, while legal, deviate from the expected stress pattern. Or, put more concretely, if IA is linked to the phonological representation of words and is, along with FA, the expected stress template in French, we anticipate that presenting words without IA impacts access to meaning and modulates the (frontal) N400.

## 2 Methods

### 2.1 Participants

20 French native speakers, aged 19 −47 (mean age 24.2), gave their written consent and volunteered to take part in the study. The study was conducted in accordance with the Declaration of Helsinki. Subjects had foreign language skills at high-school level or less, they were right-handed, with normal hearing abilities and no reported history of neurological or language-related problems. Due to excessive artifacts in the EEG signal, two participants are excluded from the EEG analyses.

### 2.2 Speech stimuli

This corpus consisted of French carrier sentences that were spoken by a native male speaker of standard French and recorded in an anechoic chamber using a digital audiotape (sampling at 44.1 kHz) (see also Magne et al., 2007). The sentences were spoken in a declarative mode, with the pitch contour always falling at the end of the sentence. Furthermore, each sentence ended with a trisyllabic target noun that either made sense in the semantic context of the sentence (semantically congruent, **+**S) or was nonsensical with its preceding context (semantically incongruent, **−**S) (see figure 1 for an example of the item **+**S and **−**S, with target words **+**IA and **−**IA). Semantically incongruous sentences were built by replacing the final congruent word with a word that shared similar acoustic and phonological characteristics, but did not make sense in the sentence context. Moreover, semantically congruent and incongruent target words all had CV syllable structures and were matched for word frequency (92.38 and 91.36 occurrences per million, respectively), using the LEXIQUE2 French lexical database (New et al., 2001, in Magne et al. 2007). So, congruent and incongruent target words were acoustically and phonologically similar and had been matched in word frequency and word and syllable duration (a more detailed account on the construction of the sentences can be found in Magne et al. 2007).

**Figure 1:**
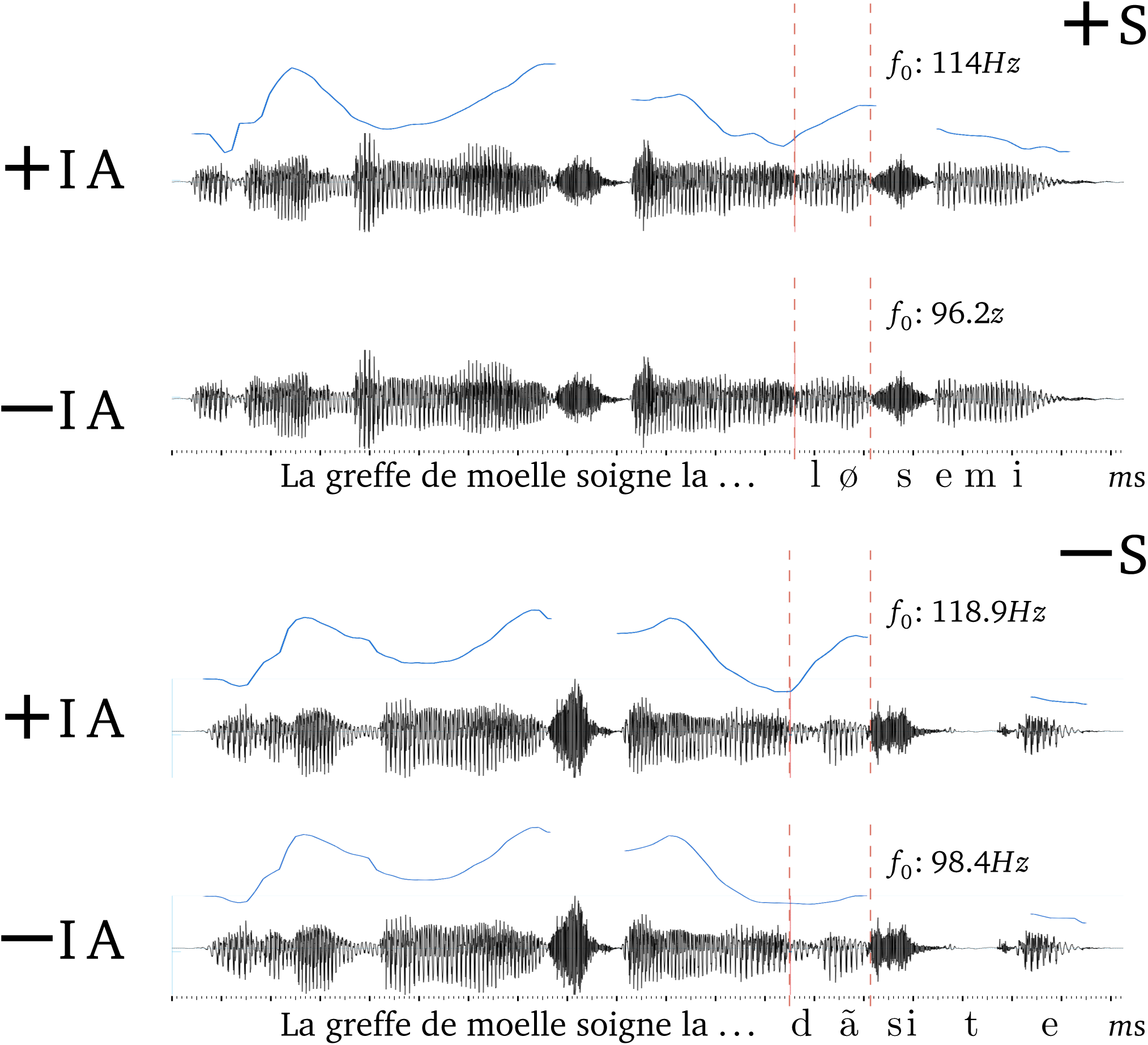
Example of *f*_0_ resynthesis with (**+**IA) and without initial accent (**−**IA) on semantically incongruent (**+**S, top two) and semantically congruent (**−**S, bottom two) sentences with quadratic interpolation from the *f*_0_ value of the preceding determinant to the *f*_0_ value at the beginning of the last stressed syllable for **+**IA targets (visible in blue). The time window of ±IA is indicated by vertical red dashed lines.

Stimuli selection was based on the presence of a marked and natural IA in the original corpus in both semantic conditions. Because the primary phonetic parameter of IA is a rise in *f*_0_ (Astésano, 2001), this meant that only sentences in which the target nouns in both semantic conditions started with a rise of *f*_0_ of at least 10% on the first syllable compared to the preceding *f*_0_ value on the (unaccented) determinant (Ladd, 2008; Astésano et al., 2007) were admitted in the current corpus. 160 stimuli met this criteria; 80 carrier sentences with 80 **+**S target nouns and 80 **−**S target nouns.

The metrical condition (± IA) was created by lowering the *f*_0_ value on the first vowel of the target-words near the *f*_0_ value on the preceding (unaccented) determinant in order to remove the natural **+**IA and create the **−**IA condition (see figure 1). This manipulation was achieved using a customized quadratic algorithm (see Aguilera et al., 2014, for more details) in PRAAT (Boersma & Weenink, 2016) which progressively modified the *f*_0_ values while allowing for micro-prosodic variations to be maintained such that the natural sound of the stimuli remained intact. Further, the **+**IA stimuli were forward and back transformed to equalize the speech quality between **+**IA and **−**IA stimuli.

The resulting 320 stimuli over the four experimental conditions (**+**S **+**IA, **−**S **+**IA, **+**S **−**IA, and **−**S **−**IA) were divided over four lists, such that each participant was presented with 80 unique sentences, i.e. 20 sentences per condition.

**Table 1:** Overview of mean stimulus properties within the semantically congruent and incongruent conditions ±IA (total sentence and target-word duration, first syllable and syllable-vowel durations, and first syllable-vowel *f*_0_ values).

| | Sentence *ms* | | Target word *ms* | | 1st syllable *ms* | | 1st vowel *ms* | | 1st vowel $f_0$ | |
| --- | --- | --- | --- | --- | --- | --- | --- | --- | --- | --- |
|  | *m* | *sd* | *m* | *sd* | *m* | *sd* | *m* | *sd* | *m* | *sd* |
| SEMANTICALLY CONGRUENT |  |  |  |  |  |  |  |  |  |  |
| -IA | 2097.07 | 402.81 | 552.88 | 96.98 | 157.23 | 28.76 | 72.16 | 25.9 | 116.56 | 11.73 |
| +IA | 2092.07 | 402.81 | 552.88 | 96.98 | 157.23 | 28.76 | 72.16 | 25.9 | 126.38 | 12.2 |
| SEMANTICALLY INCONGRUENT |  |  |  |  |  |  |  |  |  |  |
| -IA | 2122.59 | 411.72 | 583.45 | 61.72 | 160.57 | 32.16 | 77.86 | 27.23 | 123.02 | 42.18 |
| +IA | 2122.59 | 411.72 | 583.45 | 61.72 | 160.57 | 32.16 | 77.86 | 27.23 | 140.28 | 44.26 |

### 2.3 Procedure

Each participant was comfortably seated in an electrically shielded and sound attenuated room. Stimuli were presented through headphones using Python2.7 with the PyAudio library on a Windows XP 32-bit platform. Participants were instructed to judge as quickly and accurately as possible whether a sentence was semantically congruent or incongruent by pressing the left or right arrow key on a standard keyboard using their dominant, right hand. Arrow key assignment was counter-balanced across participants. The I S I was fixed at 600 ms. Participants were allowed to give their answer from the start of the target word until 1500 ms post stimulus offset. To ensure participants understood the task requirements, the experiment began with a short practice phase, consisting of 10 trials that were similar to the experimental trials, but not included in the analyses.

Each participant listened to a complete list of all 80 stimuli. Using Latin square designs, the four conditions (**+**S **+**IA, **−**S **+**IA, **+**S **−**IA, and **−**S **−**IA) were evenly distributed over two blocks, with block order balanced across participants. Total duration of the experiment, including the set-up of the EEG electrodes, was approximately 1.5h.

### 2.4 EEG recording and preprocessing

EEG data were recorded with 64 Ag/AgCl-sintered electrodes mounted on an elastic cap and located at standard left and right hemisphere positions over frontal, central, parietal, occipital and temporal areas (International 10/20 System; Jasper, 1958). The EEG signal was amplified by BioSemi amplifiers (ActiveTwo System) and digitized at 2048 Hz. The data were preprocessed using the EEGlab package (Delorme & Makeig, 2004) with the ERPlab toolbox (Luck et al., 2010) in Matlab (Mathworks, 2014). Each electrode was re-referenced offline to the algebraic average of the left and right mastoids. The data were band-pass filtered between 0.01 −30 Hz and resampled at 256 Hz.

Following a visual inspection, signal containing E M G or other artifacts not related to eye-movements or blinks was manually removed. I C A was performed on the remaining data in order to identify and subtract components containing oculomotor artifacts. Finally, data were epoched from −0.2 to 1 seconds surrounding the onset of the target word and averaged within and across participants to obtain the grand-averages for each of the four conditions (**+**S **+**IA, **+**S **−**IA, **−**S **+**IA, **−**S **−**IA).

### 2.5 Analysis—behavioral and EEG

#### 2.5.1 Behavioral

The behavioral data (i.e. accuracy rates and response latencies) were analyzed in R (Team, 2014) with the lme4 package (Bates et al., 2012). Visual inspection of residual plots did not reveal any obvious deviations from homoskedasticity or normality.

For the accuracy rates, binary logistic regression was used to analyze the two predictors semantic congruency and presence of IA. That is, the model tested how well semantic congruency and presence of IA predicted the proportion of errors. For response latency (a continuous variable), a linear mixed effects model was used to analyze the effect semantic congruency and IA had on reaction times. For both accuracy rates and response latencies, the models additionally included participants and stimuli as random variables. More specifically, for the random structure, intercepts for listeners and stimuli, as well as by-stimuli random slopes for the effects of metrical pattern and semantic congruity best accounted for underlying random variability. *p*-values were obtained by likelihood ratio tests of the model with the effect in question against the model without the effect in question.

#### 2.5.2 EEG

The EEG data was analyzed with a mass univariate permutation test, which allows for correction of multiple comparisons and rigorous control of the family-wise error rate, while remaining statistically powerful (Groppe et al., 2011; Luck, 2014; Fields, 2017). The analysis was implemented using the Mass Univariate ERP Toolbox (Groppe et al., 2011) and Factorial Mass Univariate ERP Toolbox (Fields, 2017) in Matlab (Mathworks, 2014) and statistical significance was assessed with the *F* _max_ statistic (Blair & Karniski, 1993). The null distribution was estimated by repeatedly sampling the data types from the ERP data, and selecting the largest *F*-value for each comparison (i.e. the *F* _max_). In all analyses, we set the number of permutations per comparison to 10, 000 to approximate the null distribution for the customary family-wise alpha (*α*) level of 0.05. To further maximize statistical power and reduce the number of comparisons, data were down-sampled to 128 Hz.

Because, while the N400 resulting from semantic incongruities is typically maximal in the centro-parietal region of the brain (Brown & Hagoort, 1993; Kutas & Federmeier, 2011), violations in metrical/phonological expectancies more commonly result in a N400 that is more frontally located (e.g. Böcker et al., 1999; DeLong et al., 2005; Lau et al., 2008; Steinhauer & Connolly, 2008; Rothermich et al., 2010, 2012; Yan et al., 2017), we selected fronto-central and centro-parietal electrodes (Fpz, FCz, Fz, AFz, Fp1, Fp2, FC1, FC2, F1, F2, AF3, AF4, Cz, P1, P2, C3, C4, Pz, P3, P4, CP1, CP2). Furthermore, because the phonological/metrical N400 has been reported to precede the semantic N400 temporally (e.g. Magne et al., 2007; Steinhauer & Connolly, 2008; Rothermich et al., 2010, 2012) we tested two separate time-windows; 351 −451 ms for the metrical N400 and 450 −650 ms for the semantic N400.

Finally, to make sure that modulations in our N400 time-windows would not reflect P2 residue due to differential acoustic processing on our ±IA stimuli, we also tested this time-window from 181 −281 ms.

## 3 Results

### 3.1 Behavioral data

#### 3.1.1 Response accuracy

There was a significant main effect of ±IA with participants making more errors when stimuli had been presented **−**IA than when they had been presented **+**IA (*β* = 1.58, *SE* = 0.63, *t* = 2.51, *p* < 0.05). The semantic condition was revealed a marginal predictor of error rate, with more errors when sentences were semantically congruent, than when they were semantically incongruent (*β* = 1.73, *SE* = 0.94, *t* = 1.85, *p* = 0.06). Interestingly, the error rates reported here are similar to those reported in Magne et al. (2007), with most errors on sentences that were semantically congruent, but metrically unexpected (note that the metrical manipulation actually created an *illegal* pattern in Magne et al. 2007). Presence of IA and semantic congruency did not interact (*β* = −0.3, *SE* = 1.26, *t* = −0.24, *p* = 0.81, *ns*).

#### 3.1.2 Reaction times

As can be seen in figure 2, both IA and semantic congruity affected response latencies. When stimuli had been presented **−**IA, participants were slower to respond than when they had been presented **+**IA (*β* = 21.0, *SE* = 9.37, *t* = 2.24, *p* < 0.05). Furthermore, as mentioned above, semantic congruity also affected reaction times (*β* = −78.46, *SE* = 16.81, *t* = −4.67, *p* < 0.001); congruent sentences were responded to faster than incongruent sentences. This effect was expected and is in line with the results reported in Magne et al. 2007. Presence of IA and semantic congruency did not interact (*β* = 10.66, *SE* = 18.04, *t* = 0.59, *p* = 0.55, *ns*).

**Figure 2:**
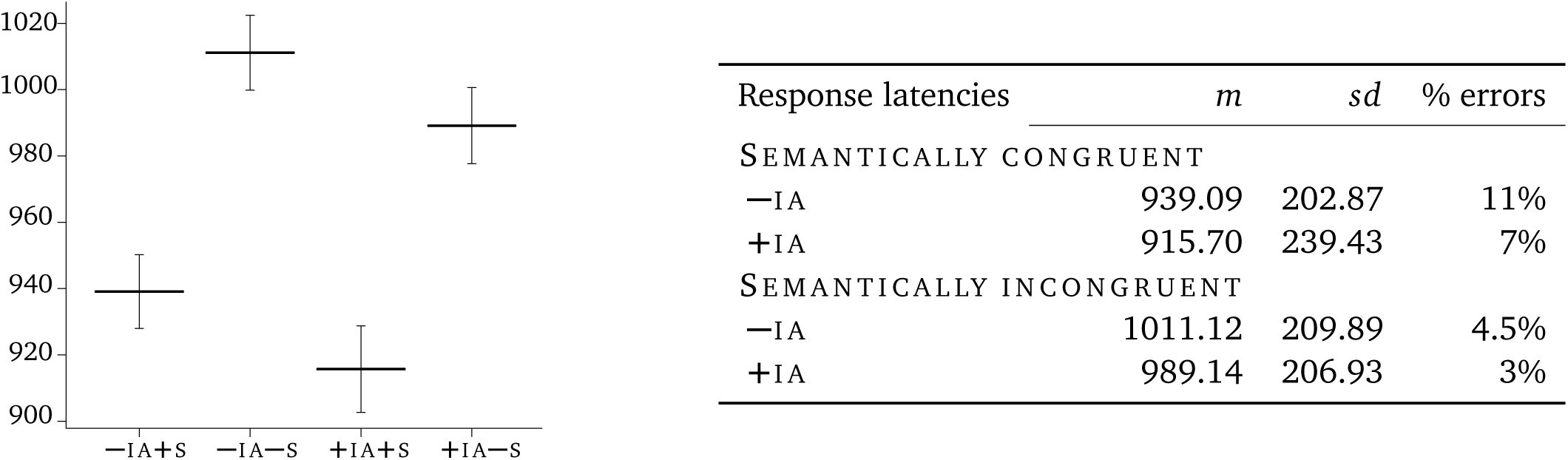
Left: Error-bar plot of mean reaction times for all four conditions (**−**IA**+**S, **−**IA **−**S, **+**IA **+**S, **+**IA **−**S) revealing a significant effect of both ±IA and of ±S, with no interaction between the two experimental manipulations. Right: Reaction times and error rates per condition. Data analysis revealed a significant effect both of ±IA and of ±S, with no interaction between the two conditions.

### 3.2 EEG

#### 3.2.1 P2

As expected, neither ±IA nor ±S modulated the P2 amplitude (*p* = 0.42 and *p* = 0.59, *ns* respectively). This means that differences we find on the later metrical N400 and semantic N400 cannot be attributed to differences on the early P2, held to reflect more bottom-up processing of purely acoustic information (Hillyard & Picton, 1987).

#### 3.2.2 Early time-window: 351 −451

The data reveal a main effect of ±IA, i.e. ±IA words modulated the metrical N400 regardless of semantic congruency (critical *F*-score: ±15.32, *d f* = 17, *p* < 0.05). Words **−**IA elicited a larger N400 than did words **+**IA 375 ms post target word onset in the anterio-frontal region (Afz) (see figure 3a).

**Figure 3:**
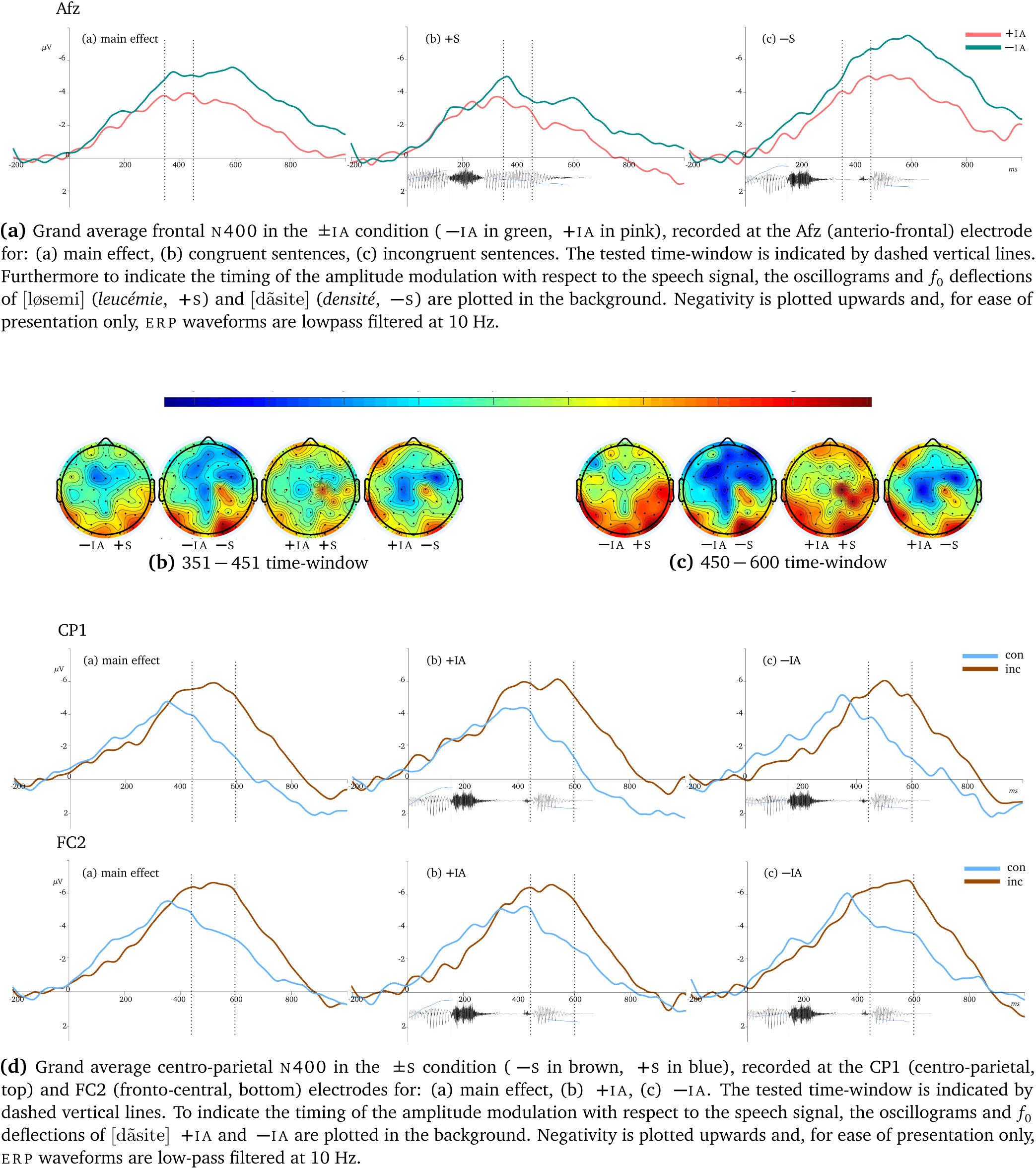
Overview of ERP data with in figure 3a the frontal N400 in the ±IA condition recorded at the Afz, in figures 3b and 3c the topographic mean amplitude maps for the early (351 −451) and late (450 −600) time-windows, respectively, and in figure 3d the centro-parietal N400 in the ±S condition recorded at the CP1.

Semantic congruency had no effect on the metrical N400 (*p* = 0.14, *ns*) nor did it interact with presence of IA (*p* = 0.15, *ns*).

#### 3.2.3 Late time-window: 450 −600

In this later time-window, the ERP data show a main effect of semantic congruity (critical *F*-score: ±17.47, *d f* = 17, *p* < 0.05): semantically incongruent sentences elicited a larger N400 between 492 −593 ms after the onset of the target word than semantically congruent sentences in the left centro-parietal region (CP1) and the right fronto-central region (FC2) (see figure 3d). This difference in N400 amplitude was also significant within the condition **−**IA (critical *F*-score: ±17.05, *d f* = 17, *p* < 0.05) and not significant within the condition **+**IA (*p* = 0.086).

The main effect of IA was not significant (*p* = 0.28, *ns*), however, we did observe an interaction between ±IA and ±S. The interaction effect between our two manipulations was significant between 523 −593 located at centro-parietal and frontal electrodes (critical *F*-score: ±15.86, *d f* = 17, *p* < 0.05, at Af4, Afz, CP1 and FC2), such that, in this later time-window, ±IA had continued to modulate N400 amplitude, but only in the sentences that were semantically incongruent ((critical *F*-score: ±17.09, *d f* = 17, *p* < 0.05, see figure 3a).

Furthermore, visual inspection suggested a difference in N400 onset latency between semantically congruent and incongruent sentences, but only in the **−**IA condition, indicating that conflict resolution starts later for incongruent words without initial accent. Because this visual effect is important for the discussion of the additional semantic processing cost when words are presented **−**IA, we computed a regression analysis with, as dependent variable, peak amplitude latency, ±IA, semantic congruency and electrode cite (parietal, centro-parietal and central) as fixed effects, and participants as random effects. However, the analysis was not significant at *p* = 0.11. These results are further interpreted below.

## 4 Discussion

In the present study, we examined the phonological status of the French initial accent and its role in semantic processing. We were particularly interested in modulations of the N400 ERP component, a component typically observed subsequent to violations of lexico-semantic expectations (e.g. Kutas & Hillyard, 1980; Brown & Hagoort, 1993). Below, we present each of our findings in turn, starting with the main effect of ±IA on the fronto-central metrical N400 to then discuss the interaction between metrical expectancy and semantic congruence on the centro-parietal N400. Finally, we will examine our behavioral data which suggest that violated metrical anticipations slow down semantic conflict resolution during speech processing.

### 4.1 Metrical N400

During the early N400 time-window, presence of initial accent modulated N400 amplitude in the anterio-frontal brain area, irrespective of semantic congruency, i.e. words without initial accent elicited a larger N400 than did words with initial accent (figure 3a). This, again, indicates that listeners expected words to be presented with initial accent, in line with the results reported in te Rietmolen et al. (2016). Furthermore, because our manipulation of IA did not modulate the acoustic P2, the metrical effect can be interpreted to reflect a more controlled process in the phonological processing of the initial accent, i.e. IA is phonologically natural.

Note also that, because in this time-window, we observed a main effect of our ±IA manipulation, this negativity may well be another instance of the N325 and indicate difficulties in stress extraction during lexical access (*cf.* Böcker et al., 1999).^2^ This finding has two important consequences for interpreting the role of IA and more generally the domain of accentuation in French. First, replicating the results reported in te Rietmolen et al. (2016) is far from trivial for a language allegedly without accent wherein stress is not lexically distinctive and has been mostly ignored by the scientific community. Replication is at the core of science, and particularly the functional value of IA—the traditionally secondary and optional accent—has been largely overlooked. Moreover, while there has been more scientific interest for the contributions of stress in speech comprehension in stress based languages, metrical stress extraction during speech processing as reflected by a modulation of the N325 had only been shown in Böcker et al. (1999), and predominantly when listeners were performing a stress discrimination task requiring them to *explicitly attend* the metrical information. Replicating the N325 effect in the current work, even when using a different paradigm, shows that French listeners have a metrical expectation for the initial accent, which they extract automatically during speech processing and use in the task at hand, i.e. lexical retrieval and semantic access.

The second conclusion we can draw in observing a main effect of IA in the current study, is that stress extraction is hindered when words are presented without their expected initial accent marking their onset, *even* when the word is *embedded* within a sentence (i.e. not presented in isolation). Indeed, as was explained above, the previous ERP studies had always manipulated IA on isolated words where the accent was in utterance initial position, which made it difficult to rule out advantages applying to the levels higher in the prosodic hierarchy. Here, however, we obtain the same effects despite IA not being utterance initial, underscoring the phonological status of IA as marker of the left boundary of the word (*cf.* Astésano et al., *2007, 2012; Garnier et al., 2016; Garnier, 2018).*

### 4.2 Semantic N400

During the later N400 time-window, semantic congruity modulated the N400 in the centro-parietal regions, with semantically incongruent sentences eliciting a more ample N400 than did semantically congruent sentences (figure 3d). This effect was however more pronounced when words were presented without IA than when they had been presented with IA, suggesting an interaction effect between semantic congruity and metrical expectation, such that pre-semantic processes (in this case the extraction of the initial accent) facilitated subsequent semantic evaluation. Indeed, arguably, the processes of word recognition and semantic retrieval unfold, due to the temporal nature of speech input, in a cascading manner. Phonological analysis would then be required before semantic evaluation and this analysis is likely facilitated when the input meets phonological and metrical expectations (see also e.g. Rothermich et al., 2010, 2012).

Note that the findings therefore indicate that speech comprehension is impaired when the analysis of unexpected metrical stress templates has a downstream impact on semantic retrieval and integration (e.g. Praamstra & Stegeman, 1993; Dumay et al., 2001; DeLong et al., 2005; Robson et al., 2017). The results then contradict the hypothesis that the N400 can only be modulated by hindered post-lexical processes such as contextual integration (van den Brink et al., 2001; Brown & Hagoort, 1993), and, instead suggest phonological processes also affect N400 amplitudes. In this view, the N400 thus reflects the degree of lexical pre-activation with higher levels of pre-activation facilitating lexico-semantic processes and reducing N400 amplitude (Kutas & Hillyard, 1980; Kutas & Federmeier, 2011; DeLong et al., 2005; Gilbert, 2014).

Such a view takes the N400 to reflect predictive, anticipatory processes that need not exclusively be of semantic nature, but can be phonological as well (Praamstra & Stegeman, 1993; Dumay et al., 2001; DeLong et al., 2005; Lau et al., 2008; Robson et al., 2017). That is, our results suggest that semantic as well as phonological predictions are generated prior to bottom-up information becoming available. Frontal regions are suggested to be involved in the generation of expected information that drive top-down modulations of sensory processing (Desimone & Duncan, 1995) and may replace missing speech information (Shahin et al., 2009; Boulenger et al., 2011). Such a ‘phonological illusion’ may account for the findings reported in Jankowski et al. (1999) where the initial accent was perceived, even when its phonetic correlates were suppressed, and may account for the ERP modulations observed in the current study. In fact, because the acoustic manipulations in Jankowski and colleagues were different than the manipulations here (i.e. they had mainly manipulated the onset duration, with *f*_0_—the modulated phonetic parameter in the current study—neutralized), the combined results further point to the metrical weight and phonetically-independent identity of the initial accent, although future (perception) studies are needed to better understand the neural mechanisms underlying the superposition of metrical stress.

Moreover, we observed an interaction effect between semantic congruity and the presence of the initial accent, such that ±IA continued to modulate N400 amplitudes, but only when sentences were semantically incongruent (see figure 3a). This suggest that when a word did not make sense in the semantic context of the sentences, listeners re-evaluated the phonological make-up of the word. So, our results support the idea that listeners have a phonological preference for words to be marked with the initial accent in their underlying stress pattern.

### 4.3 Delayed semantic resolution

Visual inspection of the ERP waveforms further suggested a delay in N400 latency (although this latency difference was not significant) when semantically incongruent words had been presented without initial accent, indicating that, when words are presented without initial accent and thus mismatch the listener’s metrical anticipation, semantic conflict resolution starts later. Our behavioral results are in line with this interpretation. The results in response latencies showed a main effect of IA, such that when words were presented without initial accent, participants were slower to respond than when they had been presented with initial accent. This, indeed, suggests that semantic ambiguities were resolved after participants had attended to the metrical hindrance when words were presented without their expected stress template.

We also obtained a main effect of ±IA on error rates, such that listeners made more errors when words had been presented without initial accent than when they had been presented with IA. Furthermore, listeners appeared to make most errors on sentences that were semantically congruent, but metrically unexpected, indicating that presenting the words without the initial accent misdirected the participants on the word’s identity. This is in line with the results reported in Magne et al. (2007), wherein metrical *congruity* was manipulated (i.e. the authors lengthened the medial syllable, a violation in French), while here we manipulated metrical *probability* (i.e. the presence of the initial accent). Whereas we predicted listeners to prefer words to be presented with initial accent, reducing its phonetic correlates did not create an illegal stress pattern. Still finding an effect of ±IA thus shows a *strong* expectation for the, allegedly, secondary and *optional* French accent.

Together with our ERP results on the semantic N400, the findings suggest a strong memory trace for the initial accent, such that lexical candidates matching the memory trace are easier to recognize, responded to faster and generate smaller N400s than when candidates are less easy to match (i.e. hold a less established memory trace). In other words, if listeners continuously predict upcoming speech input, they may have prepared for expected upcoming words by activating their expected phonological, metrical and semantic features from the mental lexicon (e.g. Lau et al., 2008). When all these features mismatched, reaction times were slowed down, and ERP amplitudes and latencies, which index prediction errors, increased.

## 5 In conclusion

In sum, we investigated the status of the French initial accent and its function in lexico-semantic processing. The initial accent was previously thought of as an optional and secondary accent in French, sub-serving the primary final accent in the marking of phrase boundaries. Previous ERP studies which also investigated the phonological status of IA (e.g. Astésano et al., 2013; Aguilera et al., 2014; te Rietmolen et al., 2016) showed a phonological expectancy for IA and a disruption in pre-lexical stress processing when IA had been omitted. However, in the studies, words were presented in isolation, with IA in utterance initial position. Therefore, it had remained unclear whether the facilitatory effects of IA really applied to the lexical domain. In the current study, the initial accent was not utterance initial but embedded in a sentence. We found the presence of IA to modulate the N400 not only in the fronto-central brain regions, but also in the centro-parietal regions. That is, when asking listeners to judge the semantic congruity of sentences that differed only in the explicit presence of the initial accent, lexico-semantic processing (as reflected by the N400) was still affected. Pre-lexical stress templates serve to access the mental lexicon. Our data demonstrate that presenting words without IA obstructs lexical access, which in turn, cascades up the process of speech comprehension to additionally hinder post-lexical processing. In other words, French speech processing naturally and automatically engages metrical stress processing.

## Acknowledgments

This study was supported by the Agence Nationale de la Recherche grant ANR-12-BSH2-0001 (PI: Corine Astésano).

## Footnotes

1 Note that while the post-lexical integration theory *may* reject anticipatory processes and consider the N400 to index exclusively post-lexical processes initiated upon perceiving the target word, it not necessarily *needs* to; one can easily imagine integration processes to also benefit from successful (semantic) anticipation based on prior contextual information (as is pointed out by Yan et al. 2017, see also Kuperberg & Jaeger 2016 and Nieuwland et al. 2018).

2 Note that the negativity could also be and instance of the previously reported frontal N400 (Dittinger et al., 2017; François et al., 2017) in which case it would reflect novel word-form to meaning mapping. However, even if, in this study, listeners were expected to prefer words to be marked with IA, words without IA are not illegal in French, i.e. in continuous speech IA is not always fully realized and may be suppressed to serve for instance a more rhythmically balancing function. So, French listeners are expected to be quite familiar with the stress templates **−**IA as well. Moreover, stress is never lexically distinctive in French (word meaning never changes depending on the location of the stress) so we consider it unlikely for French listeners to perceive the **−**IA stress templates as “new” auditory word forms, which they would be tasked to attach to established semantic representations.

